# Methionine regulates antitumor function of CD8^+^ T cells through polyamine synthesis

**DOI:** 10.1101/2025.10.03.680201

**Authors:** Tian Zhao, Gillian A. Carleton, Sarah Macpherson, Madison Shiyuk, Joseph Monaghan, Jun Han, Oro Uchenunu, Robert Rottapel, Ralph J. DeBerardinis, Kelly M. Stewart, Bo-Hyun Kim, Juan Ausió, David R. Goodlett, Helena Pětrošová, Kyle D. Duncan, Julian J. Lum

## Abstract

Methionine is an essential amino acid critical for T cell activation. While methionine restriction (MR) combined with immune checkpoint blockade has been shown to enhance T cell function, the impact of methionine on adoptive T cell therapies is largely unexplored. Here, we examined the functionality of T cells under MR and pharmaceutical inhibition of the methionine cycle (MAT2Ai), using primary T cells and a murine adoptive T cell therapy model. *In vitro*, transient MR or MAT2Ai treatment increased interferon gamma (IFNγ) expression in CD8^+^ T cells, whereas sustained MR led to the upregulation of T cell exhaustion-associated markers. Mechanistically, transient MR suppressed the polyamine synthesis pathway, and supplementation with polyamines reversed MR-induced IFNγ expression. Genetic ablation of s-adenosylmethionine decarboxylase, an enzyme in the polyamine synthesis pathway, recapitulated the effect of MR, indicating that transient MR enhances T cell function by inhibiting polyamine synthesis. Despite this, transient MR treatment of ovalbumin (OVA)-specific (OT-I) CD8^+^ T cells prior to adoptive transfer did not improve antitumor efficacy against EG7-OVA tumors *in vivo*. In contrast, sustained dietary MR accelerated EG7-OVA tumor growth in mice treated with OT-I T cells, demonstrating that methionine availability is essential for the activity of adoptively transferred T cells. These findings suggest that enhancing methionine availability in the tumor microenvironment may improve the efficacy of adoptive T cell therapies.

## Introduction

Effective tumor immunity relies on the capacity of cytotoxic CD8^+^ T cells to directly lyse malignant cells. Sustaining this effector function requires tightly coordinated biosynthetic and bioenergetic programs that fuel activation, proliferation, and cytokine production (1). Beyond this, these metabolic circuits are intrinsically linked to T cell fate decisions and effector function (1).

Among these key metabolic inputs, amino acids play dual roles as both substrates for protein synthesis and precursors for metabolites that shape the magnitude and quality of T cell responses (2). For example, branched-chain amino acids enhance glucose metabolism, synergize with anti-programmed cell death protein 1 (PD-1) therapy (3), and improve CAR-T cell efficacy (4). Elevated intracellular arginine promotes oxidative phosphorylation and drives a memory-like phenotype associated with enhanced tumor control (5). Other amino acids, such as serine and glycine, are required for effector cell proliferation via the one-carbon pathway, while serine deprivation impairs nucleotide biosynthesis (6). In contrast, tryptophan catabolism via the kynurenine pathway suppresses T cell function by inducing PD-1 and downregulating effector molecules (7). Glutamine, a central amino acid for the TCA cycle, also contributes to the biosynthesis of nucleotides, amino acids, and lipids (8). The absence of glutamine impairs mTOR signaling, blunts CD8^+^ and Th1 responses, and skews differentiation toward regulatory T cells (9). Recent reports show that leucine modulates mTORC1 activity and metabolic reprogramming (10) while cysteine maintains redox balance through glutathione synthesis (11). However, amino acid signaling is not uniformly beneficial. A recent CRISPR screen revealed that excessive mTORC1 activation via the amino acid transporters SLC7A1 and SLC38A2 can impair memory T cell formation (12). Thus, immunometabolic roles of amino acids are context-dependent and capable of enhancing and constraining antitumor immunity.

Methionine plays a distinct role as the initiating amino acid for protein synthesis and regulates epigenetic and metabolic processes through the methionine cycle (5,6). In this cycle, methionine adenosyltransferase 2A (MAT2A) converts methionine into S-adenosylmethionine (SAM), a universal methyl donor required for histone and DNA methylation that governs T cell activation, differentiation, and lineage stability (13,14). Beyond its role in epigenetic regulation, SAM can be decarboxylated by S-adenosylmethionine decarboxylase (AdoMetDC). This reaction supplies an aminopropyl group to putrescine, leading to the production of the polyamines spermidine and spermine (7), metabolites that support Th1 and Th2 differentiation (8). Notably, CD4^+^ T cells deficient in polyamine biosynthesis show increased IFNγ production (8) and improved antitumor responses (9), suggesting that polyamine synthesis suppresses T cell function.

Modulating methionine metabolism can either enhance or impair T cell function, depending on the context. In the tumor microenvironment (TME), methionine restriction reduces histone methylation and increases expression of inhibitory receptors such as programmed cell death protein 1 (PD-1) (15,16). Recent work by Bian *et al*. (17) showed that tumor cells can outcompete T cells for methionine, leading to diminished histone methylation and effector function, suggesting that methionine is crucial T cell antitumor activities. In support of this, Sharma *et al*. (18) found that methionine restriction during T cell activation promotes CD8^+^ T exhaustion by enhancing nuclear factor of activated T-cells 1 (NFAT1) signaling, which impairs T cell antitumor function. Paradoxically, systemic methionine deprivation through a methionine-restricted diet, when combined with immune checkpoint blockade (ICB), increases tumor-infiltrating lymphocytes (TIL) and upregulates granzyme B expression, suggesting that methionine availability may differentially shape immune responses depending on timing, tissue context, and therapeutic co-interventions (15-17,19-22).

In this study, we investigated the role of methionine on T cell function in the context of adoptive T cell therapies (ACT), using methionine restriction (MR) and Met cycle inhibition through a MAT2A inhibitor (MAT2Ai). *In vitro*, we observed that transient MR or short-term exposure to MAT2Ai treatment enhanced effector function of activated CD8^+^ T cells, including increased cytokine production and cytotoxic potential. However, sustained MR led to elevated expression of exhaustion-associated markers in activated CD8^+^ T cells. *In vivo*, transient MR during T cell culture did not significantly improve the antitumor efficacy of adoptively transferred T cells. However, sustained dietary methionine restriction markedly diminished tumor control, indicating that continuous methionine availability is essential for maintaining robust ACT responses. Together, these findings reveal a time-dependent role for methionine metabolism in shaping the function of T cells and suggest that timing and duration of metabolic modulation are critical considerations in ACT strategies.

## Methods

### Mice

All animal studies were performed according to Canadian Council for Animal Care guidelines and approved by the University of Victoria’s Animal Care Committee. B6 Thy 1.1 mice (Strain #00406, RRID: IMSR_JAX:000406) were purchased from the Jackson Laboratory. Breeder OT-I mice (Strain #003831, RRID: IMSR_JAX:003831) and breeder B6 mice (Strain #000664, RRID: IMSR_JAX:000664) were also obtained from the Jackson Laboratory and bred in-house. All mice were housed in groups at 22 °C with a 12/12 light-dark cycle.

### Mouse tumor models

4-month-old female B6 Thy1.1 mice were randomized based on body weights and subcutaneously implanted with 1 ×10^6^ EG7-OVA tumor cells subcutaneously on the right flank on day 0. On day 5, the mice were infused with 6.5 ×10^5^ activated CD8^+^ OT-I T cells (cultured in control medium (100 μM methionine) or MR medium (5 μM methionine) for 16 h before infusion) through the tail vein. Immediately following infusion, mice were put on either a control diet (0.86% methionine, Teklad, Cat. #TD230144) or a methionine-restricted diet (0.06% methionine, Teklad, Cat. #TD230293). The composition of each diet can be found in Supplementary Table S1. Tumor volumes were measured twice per week using a digital caliper (Fisherbrand, Cat. #06-664-16), and tumor volume was calculated using the following formula: tumor volume = length x width x width/2. Mice were euthanized when the tumor volume was greater than 1500 mm^3^ or the body weight loss was greater than 20%. Animal studies were performed with the assistance provided by the Animal Care Service (ACS) at the University of Victoria as a paid service.

### Mouse T cell culture

Primary mouse CD8^+^ T cells were isolated from the spleen of B6 or OT-I mice. Briefly, mouse splenocytes were isolated by grinding a spleen against a 40 μm cell strainer (Falcon, Cat. #C352340), placed in a 6-well plate containing phosphate-buffered saline (PBS; Gibco, 1 Cat. #0010023). Splenocytes were incubated in 1 mL ACK lysing buffer (Gibco, Cat. #A1049201) for one minute at room temperature to remove red blood cells, then washed once in PBS and filtered through a 40 μm cell strainer (Falcon, Cat. #C352340). CD8^+^ T cells were isolated from the resulting cell suspension using a negative isolation kit (StemCell Technologies, Cat. #19853) following the manufacturer’s instructions. The cells were then resuspended in RPMI medium (Hyclone, Cat. #SH30255.01) containing 10% fetal bovine serum (FBS; Hyclone, Cat. #SH30396.03HI), 50 µM β-mercaptoethanol (Sigma-Aldrich, Cat. #M3148-100ML), penicillin/streptomycin (Hyclone, Cat. #SV30010), L-glutamine (Hyclone, Cat. #SH3003401), and 50 U/mL recombinant mouse IL-2 (PeproTech, Cat. #AF-212-12-100UG). To activate mouse CD8^+^ T cells, CD3 (Biolegend, Cat. #100202) and CD28 (Invitrogen, Cat. #14-0289-82) antibodies were bound to a 96-well U-bottom plate for 2h at 37 °C. The isolated CD8^+^ cells were added to the plate at 1 ×10^6^/well and incubated in a humidified CO_2_ incubator (Thermo Fisher Scientific) at 37 °C and 5 % CO2 for 2 days.

### Human T cell culture

Human peripheral blood mononuclear cells (PBMCs) were isolated from healthy donor leukapheresis products (StemCell Technologies, Cat. #70500.2) by Ficoll gradient density centrifugation. Human CD3^+^ T cells were isolated from PBMCs using human CD3 positive isolation beads (Miltenyi Biotec, Cat. #130-050-101) and LS MACS columns (Miltenyi Biotec, Cat. #130-042-401) following the manufacturer’s instructions. Isolated human T cells were activated using human CD3/CD28 T cell activator (StemCell Technologies, Cat. #10991) and cultured in RPMI medium containing 5 % human AB serum (Sigma-Aldrich, Cat. #H4522-100ML), 50 µM β-mercaptoethanol (Sigma-Aldrich, Cat. #M3148-100ML), penicillin/streptomycin (Hyclone, Cat. #SV30010), L-glutamine (Hyclone, Cat. #SH3003401), and 300 IU/mL recombinant human IL-2 (SteriMax). T cells were incubated in a humidified CO_2_ incubator (Thermo Fisher Scientific) at 37 °C and 5 % CO_2_ for 2 days.

### Tumor cell culture

EG7-OVA cells were purchased from American Type Culture Collection (ATCC) (Cat. #CRL-2113) and cultured in ATCC-modified RPMI medium (Gibco, Cat. #A1049101) containing 10% FBS (Hyclone, Cat. #SH30396.03HI), 50 µM β-mercaptoethanol (Sigma-Aldrich, Cat. #M3148-100ML), penicillin/streptomycin (Hyclone, Cat. #SV30010), L-glutamine (Hyclone, Cat. #SH3003401), and 400μg/ mL of G418 (Gibco, Cat. #10131-035) in a humidified CO2 incubator (Thermo Fisher Scientific) at 37 °C and 5 % CO_2_. EG7-OVA-Luc cells were cultured in the same medium, with the addition of 2 μg/ mL of puromycin (InvivoGen, Cat. #ant-pr-1). Only cells at low passage numbers (<20) were used in experiments.

### *In vitro* methionine restriction

Activated mouse T cells were washed once and resuspended in methionine-free RPMI medium (Gibco, Cat. # A1451701), containing 10% FBS (Hyclone, Cat. #SH30396.03HI), 50 µM β-mercaptoethanol (Sigma-Aldrich, Cat. #M3148-100ML), penicillin/streptomycin (Hyclone, Cat. #SV30010), L-glutamine (Hyclone, Cat. #SH3003401), and 50 U/mL recombinant mouse IL-2 (PeproTech, Cat. #AF-212-12-100UG). A stock methionine solution of 100mM was made freshly by dissolving L-methionine (Sigma-Aldrich, Cat. #M9625-25G) in methionine-free RPMI medium (Gibco, Cat. # A1451701) and was added to each culture condition for a final concentration of 100 µM (control), 5 µM (MR), or 0 µM (MR).

For human T cells, activated T cells were washed once and resuspended in methionine-free RPMI medium (Gibco, Cat. # A1451701) containing 5 % human AB serum (Sigma-Aldrich, Cat. #H4522-100ML), 50 µM β-mercaptoethanol (Sigma-Aldrich, Cat. #M3148-100ML), penicillin/streptomycin (Hyclone, Cat. #SV30010), L-glutamine (Hyclone, Cat. #SH3003401), and 300 IU/mL recombinant human IL-2 (SteriMax). A stock methionine solution of 25mM was made fresh by dissolving L-methionine (Sigma-Aldrich, Cat. #M9625-25G) in methionine-free RPMI medium (Gibco, Cat. # A1451701) and was added to each culture condition for a final concentration of 25 µM (control) or 2 µM (MR).

### *In vitro* inhibition of the Met cycle

To dissect the role of the Met cycle in T cell effector function, we pharmaceutically inhibited the Met cycle in activated T cells using a novel MAT2A inhibitor, AGI-24512, which has been previously described (23,24) and was provided by Servier Pharmaceuticals (formerly Agios Pharmaceuticals, Oncology). The inhibitor was reconstituted at 10 mM in dimethyl sulfoxide (DMSO; Sigma, Cat. # 67-68-5), aliquoted, and stored at -80 °C until use.

Activated mouse T cells were resuspended in RPMI medium (Hyclone, Cat. #SH30255.01) containing 10% FBS (Hyclone, Cat. #SH30396.03HI), 50 µM β-mercaptoethanol (Sigma-Aldrich, Cat. #M3148-100ML), penicillin/streptomycin (Hyclone, Cat. #SV30010), L-glutamine (Hyclone, Cat. #SH3003401), and 50 U/mL recombinant mouse IL-2 (PeproTech, Cat. #AF-212-12-100UG). AGI-24512 was diluted to 100 µM in PBS and added to the MAT2Ai culture condition for a final concentration of 1 or 5 µM, while DMSO was added to the control condition for a final concentration of 0.1%.

For human T cells, activated T cells were resuspended in RPMI medium containing 5 % human AB serum (Sigma-Aldrich, Cat. #H4522-100ML), 50 µM β-mercaptoethanol (Sigma-Aldrich, Cat. #M3148-100ML), penicillin/streptomycin (Hyclone, Cat. #SV30010), L-glutamine (Hyclone, Cat. #SH3003401), and 300 IU/mL recombinant human IL-2 (SteriMax). AGI-24512 was diluted to 100 µM in PBS and added to the MAT2Ai culture condition for a final concentration of 5 µM, while DMSO was added to the control condition for a final concentration of 0.1%.

### Flow cytometry

Around 2 ×10^5^ to 5 ×10^5^ live cells were incubated with mouse Fc receptor blocking solution (BD Biosciences, Cat. #553141) or human Fc receptor blocking solution (Biolegend, Cat. #422303) at a 1 in 10 dilution for 10 minutes at 4°C and washed once in staining buffer made by diluting 3% FBS (Hyclone, Cat. #SH30396.03HI) in PBS (Gibco, Cat. #10010023)). Then the cells were incubated in 100 μL staining buffer containing the following antibodies against T cell surface markers at 1 in 100 dilutions for 30 minutes at 4°C: for mouse T cells, BV421 anti-mouse CD3e (BD Biosciences, Cat. #562600), BV750 anti-mouse CD4 (BD Biosciences, Cat. #747102), PerCP anti mouse CD8a (BD Biosciences, 553036), RB780 anti mouse PD-1 (BD Biosciences, Cat. #755328), and APC-H7 anti mouse TIM-3 (BD Biosciences, Cat. #567165); for human T cells, BV421 anti-human CD3e (BD Biosciences, Cat. #562426), Alexa Fluor 700 anti-human CD4 (Biolegend, Cat. #300526), PerCP anti human CD8a (Biolegend, Cat. #301010), BV650 anti human PD-1 (BD Biosciences, Cat. #564104), and BV785 anti human TIM-3 (Biolegend, Cat. #345032). The cells were then washed once in staining buffer and incubated with 100 μL eFluor 506 viability dye (Invitrogen, Cat. #65-0866-14) at a 1:100 dilution for 10 min at 4°C. Thereafter, the cells were washed twice in the staining buffer and fixed and permeabilized using Foxp3 Transcription Factor Staining Buffer Set (Invitrogen, Cat. #00-5523-00) following the manufacturer’s instructions. After fixation and permeabilization, the cells were incubated in 100 μL Permeabilization Buffer (Invitrogen, Cat. #00-5523-00) containing the following antibodies at 1 in 100 dilutions for 30 minutes at 4°C: for mouse T cells, BV605 anti-mouse IFNγ (Biolegend, Cat. #505840), Alexa Fluor 488 anti TCF-1 (BD Biosciences, Cat. #567018), and Alexa Fluor 488 anti TCF-1 (BD Biosciences, Cat. #567018); for human T cells, PE anti human IFNγ (Biolegend, Cat. #502509) and Alexa Fluor 488 anti TCF-1 (BD Biosciences, Cat. #567018). After that, the cells were washed twice in Permeabilization Buffer (Invitrogen, Cat. #00-5523-00), resuspended in PBS, and analyzed on an Aurora flow cytometer (Cytek Biosciences). Flow cytometry data were acquired using SpectroFlo (Cytek Biosciences) and analyzed using FlowJo v10.10 (BD Biosciences).

### T cell cytotoxicity assay

Activated OT-I T cells were cultured in control or MR medium for 16 hours, then added to 2 ×10^4^ EG7-OVA-Luc cells at effector to target (E:T) ratios of 2:1, 1:1, 1:2, 1:4, and 1:8 for 4 hours in a 96-well white-bottom plate in a humidified CO2 incubator (Thermo Fisher Scientific) at 37°C and 5% CO2. Each group was set up in quadruplicate. Spontaneous death wells (2 ×10^4^ EG7-OVA-Luc cells) and maximum killing wells (2 ×10^4^ EG7-OVA-Luc cells in 1% Triton-X 100 (Fisher BioReagents, Cat. #BP151-100)) were also tested in quadruplicate in the same plate. Four hours after incubation, D-Luciferin (Revvity Health Sciences, Cat. #122799) was added to each well to a final concentration of 150 μg/mL and mixed thoroughly. The plate was spun briefly, and each well was read for Luminescence (L) using a Varioskan LUX Multimode Microplate Reader (Thermo Fisher Scientific). The percentage of killing was determined using the formula: % killing = [L (spontaneous death) - L (experimental killing)] / [L (spontaneous death) - L (maximum killing)] × 100.

### CRISPR-Cas9 knockout of *Amd1*

Two single guide RNAs (sgRNAs) targeting *Amd1* were designed using CHOPCHOP. The specificity of the top-rated sgRNAs was examined by mapping them to the mouse genome using the BLAT search tool on the University of California Santa Cruz (UCSC) Genome Browser. Two highly specific sgRNAs targeting exon 1 of *Amd1* were selected for experiments. sgRNA1 (5’-TCTTCGTACCATCCCAAGGT-3’) and sgRNA2 (5’-ATCTTCGTACCATCCCAAGG-3’) were synthesized by Integrated DNA Technologies (IDT).

Gene knockout was performed as follows. Briefly, 300 pmol of each sgRNA was mixed with 120 pmol Cas9 protein (Invitrogen, Cat. #A36408) and incubated at room temperature for 15 min to form ribonucleoprotein (RNP) complexes. Meanwhile, 2.5 ×10^6^ activated CD8^+^ T cells were washed once in PBS (Gibco, Cat. #10010023) and resuspended in 20 μL Lonza P4 buffer (Lonza, Cat. #V4XP-4032). The cell suspension was then mixed with the RNP complexes or an equal volume of PBS, along with 125 pmol Alt-R Cas9 Electroporation Enhancer (IDT, Cat. #1075916). The resulting mixture was transferred to a 16-well Nucleocuvette Strip (Lonza, Cat. #V4XP-4032) and electroporated in a 4D-Nuceofector (Lonza) using pulse code CM137. After electroporation, the cells were allowed to recover in a humidified CO_2_ incubator (Thermo Fisher Scientific) for 5 min in pre-warmed RPMI complete medium containing 10% FBS (Hyclone, Cat. #SH30396.03HI), 50 µM β-mercaptoethanol (Sigma-Aldrich, Cat. #M3148-100ML), penicillin/streptomycin (Hyclone, Cat. #SV30010), L-glutamine (Hyclone, Cat. #SH3003401), and 50 U/mL recombinant mouse IL-2 (PeproTech, Cat. #AF-212-12-100UG). After incubation, the cells were transferred to a 24-well plate containing warm RPMI complete medium and cultured in a humidified CO_2_ incubator (Thermo Fisher Scientific).

### Quantification of indels using TIDE analysis

Four days after electroporation, genomic DNA (gDNA) was extracted from T cells using Lucigen QuickExtract DNA Extraction Solution 1.0 (BioSearch Technologies, Cat. #SS000035-D1) following the manufacturer’s instructions. Forward and reverse primers were designed using a web tool CRISPOR (25) to amplify a DNA sequence of around 700 base pairs surrounding the cut sites. The forward primer (5’-GGTTACACAGCCCAAGGTCA-3’) and the reverse primer (5’-GCTCACGAAAATAGCCGGGA-3’) were synthesized by IDT. For PCR amplification, 100 ng gDNA was mixed with 1.8 pmol of each primer, 30 μL KAPA HiFi HotStart ReadyMix (Roche, Cat. #09420398001), and 24 μL DNase/RNase-free distilled water (Invitrogen, Cat. #10977015) in a total volume of 60 μL. The mixture was then placed in a SimpliAmp thermal cycler (Thermo Fisher Scientific) and amplified using the following program: 98°C for 3 min, 28 cycles of 98°C for 30 s, 67°C for 15 s, and 72°C for 30 s, and 72°C for 1 min. The PCR products were sent to Genome Quebec for desalting and bi-directional Sanger sequencing as a paid service. The sequences were analyzed for the presence of insertions and deletions of nucleotides (indels) in the PCR-amplified sequences using Tracking of Indels by Decomposition (TIDE) analysis (26).

### Intracellular metabolite profiling using targeted LC-MS/MS

Activated T cells were washed once in ice-cold 0.9% saline (made in-house from sodium chloride (Sigma-Aldrich, Cat. #S9625-500G) and HPLC grade water (Sigma-Aldrich, Cat. #270733-1L)), lysed in 80% methanol (made in-house from methanol (Sigma-Aldrich, Cat. #34860-1L-R) and 20% HPLC grade water (Sigma-Aldrich, Cat. #270733-1L)), and flash frozen in liquid nitrogen. The samples were further processed and analyzed by liquid chromatography-tandem mass spectrometry (LC-MS/MS) as described previously (27). Chromatogram review and peak area integration were performed using TraceFinder software version 5.1 (Thermo Scientific). Peak areas were normalized to the total ion count (TIC) for each sample, and the normalized data were log-transformed and auto-scaled using MetaboAnalyst 6.0 (28).

### LC-MRM/MS assay to determine the absolute concentration of methionine in mouse serum

Mouse blood samples were allowed to clot at room temperature, then centrifuged at 2000 x g for 10 min at 4 °C to isolate serum. The serum samples were analyzed by the University of Victoria Genome BC Proteomics Centre as a paid service. Briefly, an internal standard solution of L-methionine-(methyl-d3) (Sigma-Aldrich, Cat. #300616) at 0.01 nmol/mL was prepared in 90% acetonitrile (LC-MS grade, Fisher Scientific, Cat. #A9554). A stock solution of L-methionine hydrochloride (100 mM in 0.1 M HCl, Sigma-Aldrich, Cat. #50272) was serially diluted in a ratio of 1 to 4 (v/v) with LC-MS grade water (Fisher Scientific, Cat. #W6500) to make nine calibration solutions. The concentrations of methionine in the calibration solutions ranged from 0.01 to 2000 nM. Mouse serum samples were thawed on ice and vortexed. 20 µL of each serum sample and 20 µL of each calibration solution were aliquoted into 1.5-mL Eppendorf tubes. 80 µL internal standard solution was precisely added to each tube. The mixtures were vortex-mixed for 30s at 3,000 rpm and subsequently ultrasonicated in an ice-water bath for 1 min, which was followed by centrifugal clarification at 21,000 x g and 10 °C for 5 min. 20 µL of the clear supernatant of each solution was diluted with 180 µL of water in an LC injection micro-vial. 5 µL aliquots of the resultant sample and calibration solutions were injected into an HSS T3 UPLC column (2.1 × 100 mm, 1.8 µm; Waters) to run LC-multiple-reaction monitoring (MRM)/MS on an LC-MS/MS system. The LC-MS/MS system comprised a 1290 Infinity II UHPLC instrument (Agilent Technologies) connected to a 6495C triple-quadrupole mass spectrometer (Agilent) via an atmospheric pressure electrospray ion source (Jet Stream). The mass spectrometer was operated with positive-ion detection. The mobile phase for chromatographic separation was composed of 2 mM heptafluorobutyric acid (TCI America, Cat. #H0024) in water (solvent A) and methanol (Fisher Scientific, Cat. #AA47192M6) (solvent B) as the binary solvents for a gradient elution at 2% to 15% solvent B over 10 min, which was followed by a 4-min column equilibration between injections at 0.25 mL/min and 40 °C. The LC-MRM/MS data files were recorded and processed using MassHunter software suite (Agilent Technologies) for peak integration. To calculate the concentrations of methionine detected in mouse serum, a linear regression curve (R^2^=0.9998) of methionine within a concentration range of 1 nM to 400 nM was constructed with internal standard calibration using the data acquired from the calibration solutions. The absolute concentrations of methionine in serum were calculated by inter-plotting the calibration curve with the analyte-to-internal-standard peak area ratios measured from the sample solutions.

### Tumor embedding and sectioning

EG7-OVA tumors were embedded in Epredia M-1 Embedding Matrix (Epredia, Cat. #1310) and frozen immediately in the vapor phase of liquid nitrogen. Tissues were serially sectioned at 10 µm thickness using a Leica CM1950 cryostat (Leica) at -15°C and -10 °C for the chamber and specimen head, respectively. Sections were thaw-mounted onto polylysine-coated glass ITO slides (Poly-L-lysine, Sigma-Aldrich, Cat. #P8920; ITO slides, Delta Technologies, Cat. #CB-901N-S111) and dried for 45 minutes in a nitrogen desiccation chamber before storage at -80°C. Three serial sections were acquired from each tumor to be used for H&E staining, MALDI-MSI, and nano-DESI MSI.

### Matrix-Assisted Laser Desorption/Ionization (MALDI) mass spectrometry imaging

Tissues were removed from a -80°C freezer and equilibrated to room temperature for 45 minutes in a nitrogen desiccation chamber. On-tissue derivatization with N, N, N-trimethylammonioanilyl hydroxysuccinimidyl carbamate iodide (TAHS, Toronto Research Chemicals, Cat. #T210505) was performed as previously described, with minor modifications (29). Briefly, a 1.25 mg/mL solution of TAHS in 100% acetonitrile (can, Fisher Chemical, Cat. #AA47138M6) was applied using an HTX M5 pneumatic sprayer (HTX Technologies LLC) using the parameters described in Supplementary Table S2. Samples were then incubated in a humid chamber at 55°C for 24 hours before a 15 mg/mL solution of 2,5-dihydroxybenzoic acid (2,5-DHB, Sigma-Aldrich, Cat. #149357) in 90% ACN + 0.1% trifluoroacetic acid (TFA, Sigma Aldrich, Cat. #T6508) was applied to the tissue using an HTX M5 sprayer (HTX Technologies LLC) set to the parameters listed in Table S3.

All tissues were imaged using a timsTOF flex MALDI-2 instrument (Bruker) in positive polarity in the mass range *m/z* 80-1000. Custom laser settings with SmartBeam (Bruker) were used: 50 µm step size, 500 shots per pixel, 10,000 Hz frequency, and 10.0 µs post-ionization delay. The following settings were tuned to maximize transmission in the low mass range: funnel RF amplitudes (funnel 1: 125.0 Vpp, funnel 2: 200.0 Vpp, multipole: 200 Vpp), collision cell voltage (10.0 eV), collision RF (700 Vpp), quadrupole ion energy (5.0 eV), and pre-TOF focus (65.0 µs transfer time; 6.0 µs pre-pulse storage). External calibration was performed using an ESI-L low concentration tuning mix (Agilent Technologies, Cat. # G1969-85000) before data acquisition. Online calibration was performed during acquisition using DHB matrix ions (2,5-DHB, Sigma-Aldrich, Cat. # 149357) for reference: *m/z* 137.0223, *m/z* 155.0339, *m/z* 273.0394, and *m/z* 409.0554. Data were analyzed using SCiLS Lab software (Bruker, version 13.00.16899). Spectra were normalized to total ion current (TIC). Images were generated with a 99% hotspot quantile value.

Annotation of methionine was confirmed using on-tissue MS/MS and a methionine chemical standard (Sigma Aldrich, Cat# M9625). After MALDI-MSI, on-tissue MS/MS was performed by isolating *m/z* 326.154, representing the TAHS derivative of methionine. Collision energy was increased to 25 eV to generate fragmentation spectra from each tissue. On-tissue MS/MS spectra were compared to MS/MS spectra from a derivatized methionine standard (Supplementary Fig. S6). Control regions containing matrix and CMC embedding media were acquired beside each tissue to rule out false annotations due to chemical noise.

### Nanospray desorption electrospray ionization (nano-DESI) mass spectrometry imaging

nano-DESI mass spectrometry imaging was performed on an Orbitrap Exploris 120 using a custom interface described previously (30,31). Briefly, two fused silica capillaries (50 µm ID, 150 µm OD; Molex, Lisle, IL, USA) were positioned next to one another at an ∼90° angle. A 0.5 µL/min flow of spray solvent (9:1 MeOH [FisherScientific, Cat# A456-4]: H2O [FisherScientific, Cat# W6-4] with 0.1% formic acid [FisherScientific, Cat# A118P-500]) was delivered to the primary capillary, which was aspirated through the secondary capillary using a 2 – 3 L/min flow of nitrogen controlled using a mass flow controller (SFC-6000D-5LSPM; Sensirion). For imaging, the capillary assembly was held fixed, and the tissue sample was rastered underneath using a motorized XYZ stage (Zaber Technologies). Images were collected at a scan speed of 40 µm/sec (X direction) with 150 µm spacing between scan lines. The stages and mass flow controller were controlled using newly developed imaging software written in Python 3.12. At the MS, a capillary voltage of +3.4 kV (positive mode) was used. Full scan data was collected between *m/z* 70 – 500 at a target resolution of 120,000 at *m/z* 200.

Raw data files were aligned into pixel grids and converted to imzML format using imzML Writer (v1.1.4) (32). Regions of interest for tissue-associated pixels were manually drawn in imzML Scout (v1.1.4), and ion images were extracted using the bulk export feature (mass tolerance: 3 ppm). Violin plots were generated by pooling all pixels from the control and methionine-restricted diet groups and plotting using the matplotlib’s (3.10.3) ‘violinplot’ function. Source code is available in the Supplementary Information.

After nano-DESI MSI, the presence of methionine in the tissue was verified with on-tissue MS/MS. MS/MS data was collected for methionine under the same ion source settings as used for MSI, with a precursor m/z window of 0.5 and HCD energy of 30%. The average spectra were calculated across a line scan using the vendor software and compared with that of a 40 µM methionine standard solution (Sigma, Cat #: M-9625) measured by LC-MS/MS (Vanquish UHPLC system coupled to a Orbitrap Exploris 120, Thermo Fisher Scientific). The column was a Hypersil GOLD PEI HILIC column (150 × 1 mm; 3 µm particle size; Part # 26503-151030; Thermo Scientific) operated using acetonitrile (A; Fisher Scientific, Cat #: A996-4) and 20 mM ammonium formate aqueous solution (B; ammonium formate: Fluka, Cat #: 55674; water: Fisher Scientific, Cat #: W6-4) for elution. Initial column conditions were 90% A, which was linearly decreased to 20% over 24 mins. The gradient was then returned to 90% A over 6 mins and equilibrated for an additional 7 mins before the next injection (37 mins injection-to-injection time).

### H&E staining

Serial sections adjacent to MALDI and nano-DESI scanned sections were fixed with 10% neutral buffered formalin (Sigma-Aldrich, Cat. # HT501128) and re-hydrated in a series of ethanol (Greenfield Global, Cat. # P016EAAN) washes of the following concentrations: 100%, 100%, 90%, 70%, 50%. Slides were washed in water before staining with Gill’s Hematoxylin (Sigma-Aldrich, Cat. # GHS216). Excess stain was removed with dips in water, and tissues were blued with Scott’s Modified Tap Water Substitute (2.38 mM sodium bicarbonate [Sigma-Aldrich, Cat. # S2127], 16.62 mM magnesium sulfate [Sigma-Aldrich, Cat. # M65-500]) before staining in Eosin Working Solution (0.025% w/v eosin Y [Sigma-Aldrich, Cat. # 318906], 0.5% acetic acid [Fisher Chemical, Cat. # A11350] in 80% ethanol [Greenfield Global, Cat. # P016EAAN]). Slides were dehydrated in a series of ethanol washes with increasing ethanol concentrations before a final wash in xylenes (Sigma-Aldrich, Cat. # 247642). Cover slips were mounted using Permount medium (Fisher Chemical, Cat. # SP15-100). Finally, H&E images were acquired using an Aperio Versa scanner (Leica Biosystems).

### Statistical analysis

Statistical analysis was performed using GraphPad Prism v10.1.2 (GraphPad Software). Unpaired Student’s t-tests were performed when comparing the means of two treatment groups. One-way ANOVA tests followed by Holm-Šídák tests were performed when comparing the means of more than two treatment groups. The statistical rating system is as follows: ^∗^p ≤ 0.05; ^∗∗^p ≤ 0.01; ^∗∗∗^p ≤ 0.001; ^∗∗∗∗^p ≤ 0.0001; ^∗∗∗∗∗^p ≤ 0.00001.

### Data availability

Raw metabolome data of transient-MR-treated and control CD8^+^ T cells were generated at the Howard Hughes Medical Institute. Derived data and any remaining information are available from the corresponding author upon request. MALDI-MSI datasets are available on METASPACE (https://metaspace2020.org/api_auth/review?prj=852fdcc2-91c3-11f0-a049-e3bcfe0c9b7b&token=N746_NwJcr5P).

## Results

### Transient MR enhances interferon-gamma (IFNγ) production and cytotoxicity of activated CD8^+^ T cells

To examine the role of methionine availability on CD8^+^ T cell function, activated CD8^+^ T cells were cultured for 40 hours in methionine-restricted (MR) medium. Compared to cells maintained in methionine-replete conditions, transient MR culture led to an almost twofold increase in IFNγ expression (**Fig. 1A**). Inhibition of methionine conversion to SAM using a MAT2A inhibitor (MAT2Ai) produced a similar increase in IFNγ levels to an extent comparable to MR conditions (**Fig. 1B**). To determine whether elevated IFNγ corresponds with enhanced cytotoxicity, activated OVA-specific CD8^+^ T cells (OT-I) were preconditioned for 16 hours in MR or control medium and then co-cultured with OVA-expressing EG7 tumor cells. Transient-MR-conditioned OT-I cells exhibited greater cytolytic activity compared to control cells, particularly at higher effector-to-target (E:T) ratios (**Fig. 1C**), suggesting that transient MR enhances cytotoxicity of CD8^+^ T cells.

**Figure 1.**
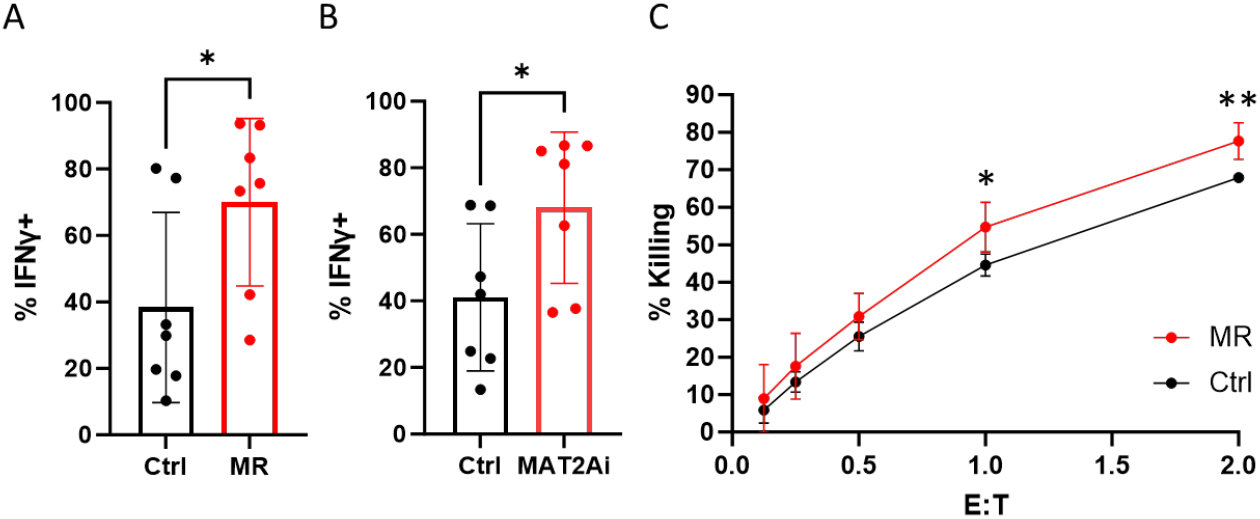
Transient MR enhances CD8^+^ T cell effector function. **A**, IFNγ expression in activated mouse CD8^+^ T cells cultured for 40 hours in methionine-restricted (MR; no methionine) or control medium (Ctrl; 100 μM methionine), assessed by flow cytometry (n=7, biological replicates). **B**, IFNγ expression in activated mouse CD8^+^ T cells treated with MAT2A inhibitor (MAT2Ai; 5 μM AGI-24512) or vehicle control (Ctrl; 0.1% DMSO) for 40 hours (n=7, biological replicates). **C**, Cytotoxicity of activated OT-I cells against EG7-OVA-Luc tumor cells following a 16-hour preconditioning in MR medium (5 μM methionine) or control medium (100 μM methionine) (n=4, technical replicates, representative of three independent experiments). Statistical significance was determined using unpaired Student’s t-tests; *p ≤ 0.05, **p ≤ 0.01. Error bars indicate standard deviation (SD).

### IFNγ expression induced by transient MR occurs via inhibition of polyamine synthesis

Methionine metabolism supports the biosynthesis of metabolites involved in redox balance, methylation, DNA synthesis, and protein translation. To investigate whether transient MR enhances T cell function through reprogramming of cellular metabolism, we performed metabolomic profiling of activated mouse CD8^+^ T cells cultured transiently in MR or control medium. This analysis revealed significant widespread changes across multiple metabolic pathways (**Fig. 2A and 2B**). As expected, intracellular levels of methionine and S-adenosylmethionine (SAM) were markedly reduced, consistent with cysteine and methionine metabolism identified as the most significantly affected pathway (**Fig. 2B**). Arginine and proline metabolism emerged as the second most impacted pathway in MR-treated cells (**Fig. 2B**). Within this pathway, SAM, the polyamines spermidine (SPD) and spermine (SPM), and 5′-methylthioadenosine (MTA) were decreased (**Fig. 2C-G**), suggesting that alterations in the polyamine synthesis pathway may be regulated by changes in methionine availability. To test whether reduced polyamine levels mediate the enhanced IFNγ expression observed under transient MR, we supplemented transient MR cultures with exogenous polyamines. Addition of SPM during transient MR reduced IFNγ expression to control levels (**Fig. 2H**). Likewise, SPD supplementation reversed the increase in IFNγ induced by MAT2A inhibition (**Fig. 2I**).

**Figure 2.**
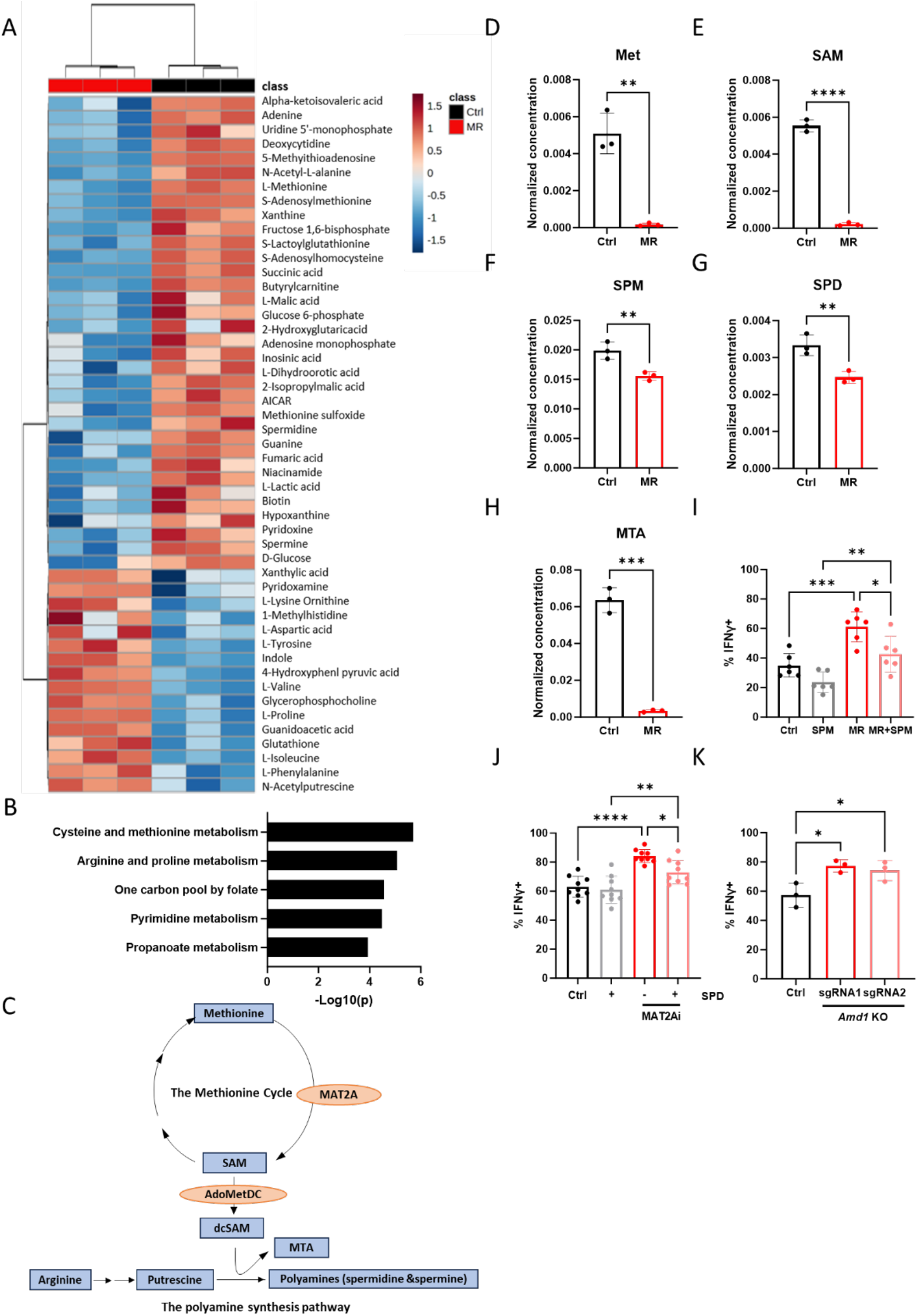
Transient MR enhances the function of CD8^+^ T cells through the inhibition of the polyamine synthesis pathway. **A**, Heat map showing the top 50 differentially abundant metabolites in activated mouse CD8^+^ T cells cultured in methionine-restricted (MR; 5 μM methionine) versus control medium (Ctrl; 100 μM methionine) for 40 hours (n=3, biological replicates). **B**, Pathway enrichment analysis reveals the most affected pathways under MR medium (5 μM methionine) for 40 h compared with those cultured in control medium (Ctrl, 100 μM methionine) for 40 hours (n=3, biological replicates). **C**, Schematic diagram showing the methionine cycle and the polyamine synthesis pathway. **D-H**, TIC-normalized concentration of metabolites in CD8^+^ T cells cultured under MR medium (5 μM Met) for 40 hours compared with those cultured in control medium (Ctrl, 100 μM Met). **I**, IFNγ expression in activated CD8^+^ T cells cultured under the following conditions: Ctrl (100 μM methionine), MR (5 μM methionine), SPM (30 μM spermine), or MR + SPM (5 μM methionine + 30 μM spermine) (n=6). **J**, IFNγ expression in CD8^+^ T cells treated with MAT2A inhibitor (1 μM AGI-24512) with or without spermidine (SPD; 50 μM); vehicle control: 0.1% DMSO (n=9, biological replicates). **K**, IFNγ expression in CD8^+^ T cells electroporated with PBS (Ctrl) or CRISPR-Cas9 ribonucleoproteins targeting *Amd1* (sgRNA1, sgRNA2) (n=3, biological replicates). Unpaired Student’s t-test (**D-H**) or one-way ANOVA with Holm-Šídák tests (**I-K**) were performed to determine statistical significance. *p ≤ 0.05, **p ≤ 0.01, ***p ≤ 0.001, ****p ≤ 0.0001. Error bars indicate standard deviation (SD).

Polyamine biosynthesis depends on the decarboxylation of SAM via adenosylmethionine decarboxylase (AdoMetDC), encoded by *Amd1*. To further examine the role of this polyamine biosynthesis on IFNγ production, *Amd1* expression was disrupted in CD8^+^ T cells using CRISPR-Cas9 with two sgRNAs targeting exon 1 of *Amd1*. Electroporation of either sgRNA efficiently knocked out *Amd1*, as confirmed by TIDE analysis, yielding indel frequencies of 63.9% and 83.9% (Supplementary Fig. S1A and S1B). *Amd1* knockout mimicked the effects of transient MR, leading to increased IFNγ production (**Fig. 2J**). Similarly, pharmacologic inhibition of AdoMetDC with CGP-48664 also elevated IFNγ levels, an effect reversed by SPD supplementation (Supplementary Fig. S1C). Taken together, these findings reveal a mechanistic link between methionine metabolism and T cell effector function, and demonstrate that transient MR enhances IFNγ production via inhibition of polyamine biosynthesis.

### Sustained MR induces exhaustion-associated markers in activated CD8^+^ T cells

While we observed a clear beneficial effect of transient MR, limited methionine availability in the TME could subject T cells to sustained, rather than transient MR. As such, the functional and phenotypic consequences of sustained MR were examined *in vitro*. Activated mouse and human T cells were cultured in MR medium for four days and assessed for IFNγ expression and markers of exhaustion. In mouse CD8^+^ T cells, sustained MR led to increased IFNγ production (**Fig. 3A**) but also elevated expression of exhaustion-associated markers, leading to increased frequencies of PD-1^+^/TIM-3^+^ and TCF-1^-^/PD-1^+^subsets (**Fig. 3B and 3C**). In contrast, sustained MR in human CD8^+^ and CD4^+^ T cells resulted in a slight lowering of IFNγ expression (**Fig. 3D and 3E**), indicating a species-specific difference in response to methionine availability. However, consistent with the mouse data, sustained MR increased PD-1^+^/TIM-3^+^ expression in human CD8^+^ T cells (**Fig. 3F**), and a similar trend was observed in CD4^+^ T cells (**Fig. 3G**).

**Figure 3.**
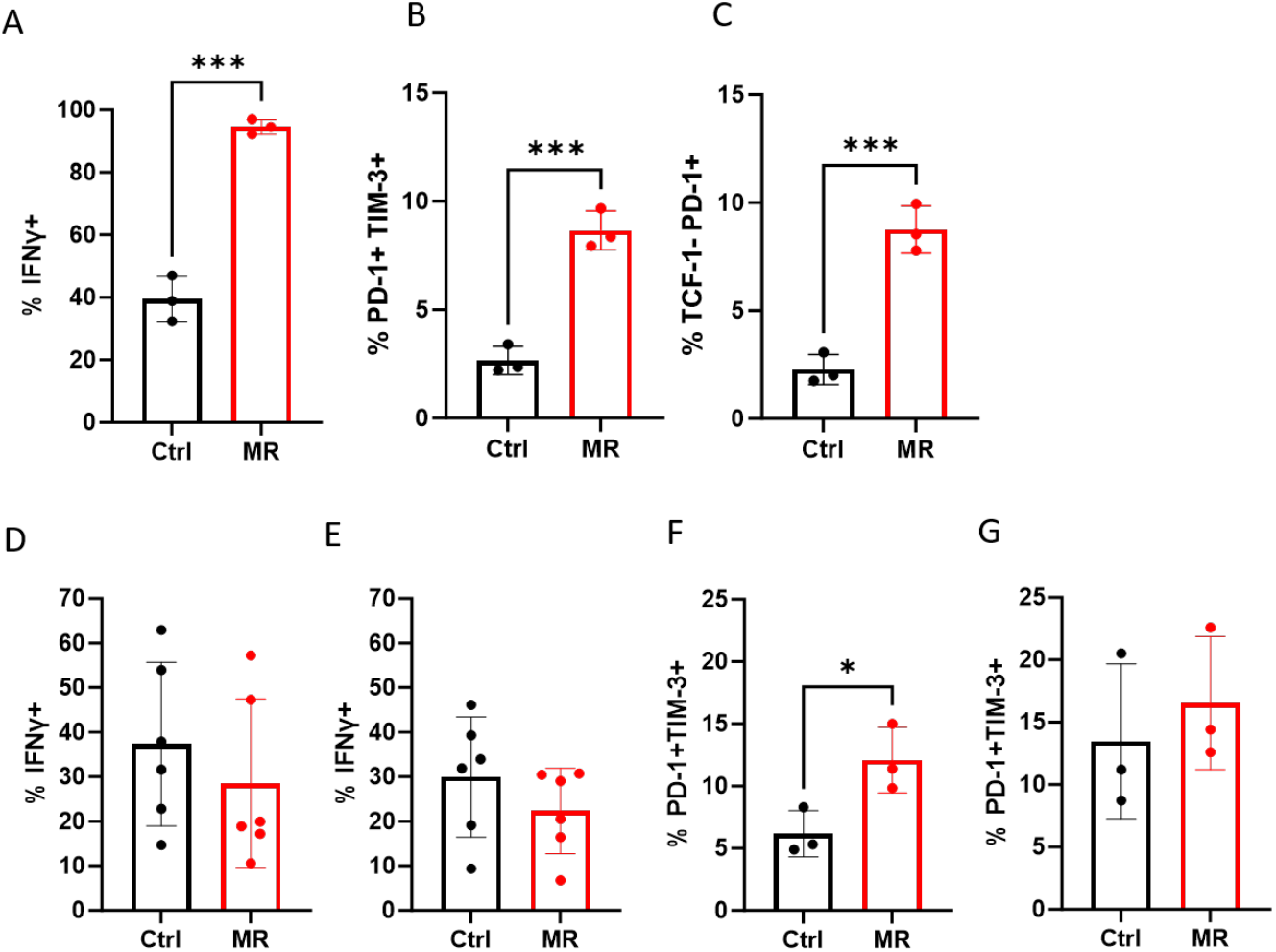
Sustained MR induces exhaustion-associated markers in CD8^+^ T cells. **A-C**, IFNγ expression and exhaustion-associated marker profiles (PD-1^+^/TIM-3^+^ and TCF-1^−^/PD-1^+^) in activated mouse CD8^+^ T cells cultured for four days in methionine-restricted (MR; 5 μM methionine) or control medium (Ctrl; 100 μM methionine), assessed by flow cytometry (n=3, biological replicates). **D-G**, IFNγ expression and PD-1^+^/TIM-3^+^ co-expression in human CD8^+^ T cells (**D and F**) and CD4^+^ T cells (**E and G**) cultured in MR medium (2 μM methionine) or control medium (Ctrl, 25 μM methionine) for 4-5 days (D and E: n=6; F and G: n=3, biological replicates). Statistical significance was determined using unpaired Student’s t-tests. *p ≤ 0.05, ***p ≤ 0.001. Error bars indicate standard deviation (SD).

To further understand how sustained MR induces exhaustion-associated markers, activated human T cells were treated for five days with the MAT2Ai. Sustained inhibition partially recapitulated the effects of MR, resulting in a trend towards decreased IFNγ expression (Supplementary Fig. S2A and S2B) in both CD8^+^ and CD4^+^ T cells and a trend towards elevated levels of exhaustion-associated markers in CD4^+^ T cells (Supplementary Fig. S2C and S2D). These findings suggest that sustained methionine deprivation induces expression of exhaustion-associated markers and that may compromise long-term antitumor immunity.

### Methionine is required for the antitumor function of adoptively transferred CD8^+^ T cells

Recent work has shown an indispensable role for methionine in early T cell activation (18). However, much less is known about how dietary methionine availability supports the durable responses of ACT, where adoptively transferred T cells are activated. To address this, the impact of transient and sustained MR was evaluated in the context of adoptive transfer of activated OT-I CD8^+^ T cells in the EG7-OVA tumor model. To achieve sustained MR in this model, we designed a standard control diet (CD, 0.86% methionine) or a methionine-restricted diet (MRD, 0.06% methionine). We first examined whether dietary MR could reduce intratumoral methionine levels. EG7-OVA-bearing mice were maintained on CD or MRD for two days after being treated with ACT. Strikingly, dietary MR significantly reduced methionine levels in tumors, as confirmed by MALDI and nano-DESI mass spectrometry imaging (**Fig. 4A and 4B**, Supplementary Fig. S3). Pooled analysis of normalized methionine intensities showed a substantial reduction in methionine levels in tumors maintained on MRD (**Fig. 4C**). This data indicate that a short course of two-day dietary MR effectively reduces intratumoral methionine levels.

**Figure 4.**
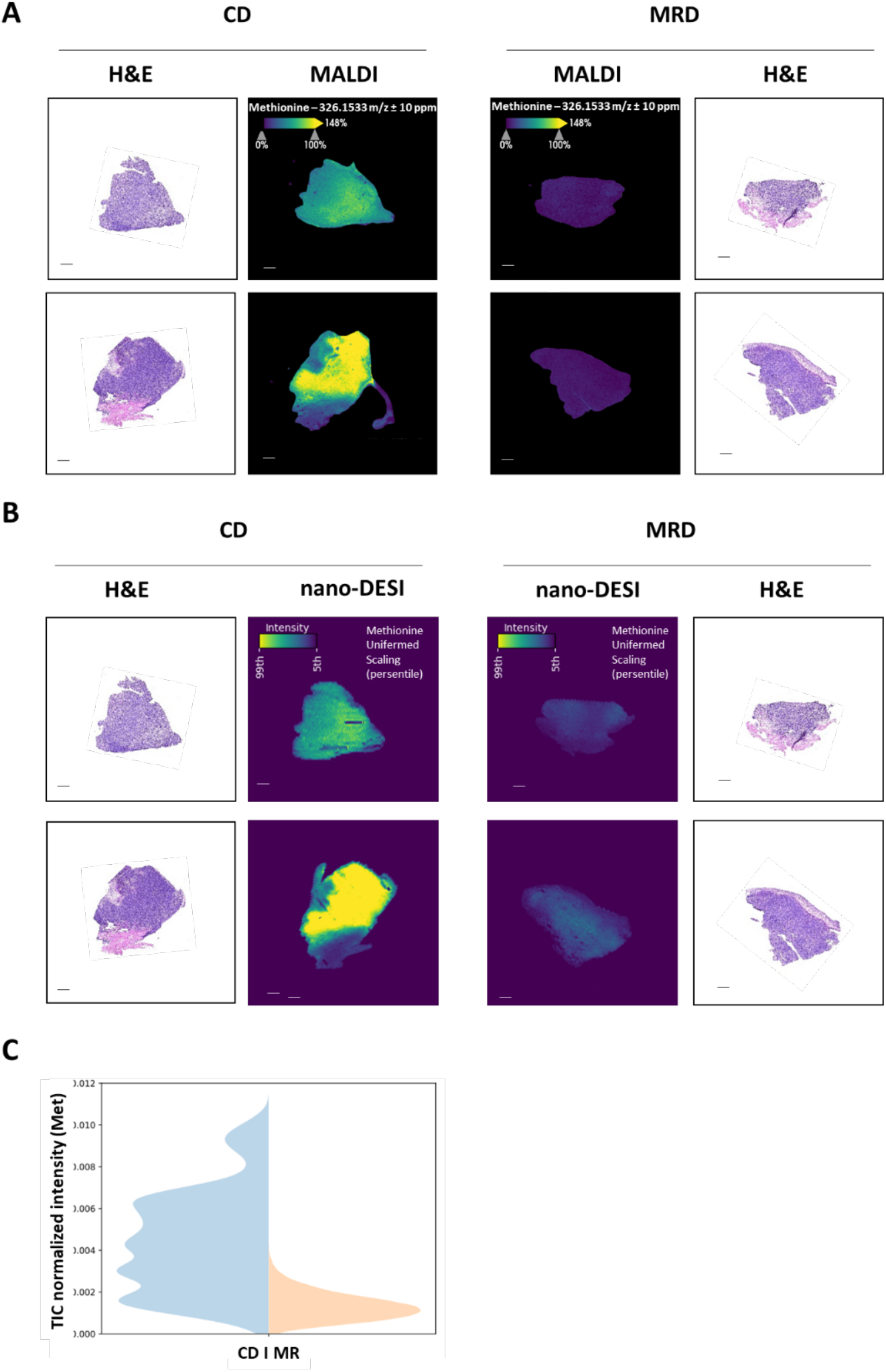
Methionine-restricted diet reduces intratumoral methionine levels. **A-C**, Female B6 Thy1.1 mice bearing EG7-OVA tumors were treated with adoptive transfer of OT-I T cells and then placed on either a control diet (CD) or a methionine-restricted diet (MRD) for 2 days. The tumors were sectioned and imaged by histochemistry microscopy (H&E staining) and MALDI or nano-DESI mass spectrometry imaging. MALDI and nano-DESI images show TIC-normalized signal intensities of methionine. **A and B**, Two representative tumors were shown for each group, scans of all tumors are shown in Supplementary Fig. S3. **A**, Representative MALDI images of tumor sections from mice placed on CD or MRD (n=6, biological replicates). **B**, Representative nano-DESI images of tumors from mice placed on CD or MRD (n=6, biological replicate). **C**, Violin plot of pooled TIC-normalized signal intensities of methionine (Met) in tumors (combined data from scans of 6 tumors per group) imaged by nano-DESI. Scale bars indicate a length of 1 mm.

We then investigated the role of transient and sustained MR in regulating antitumor function of adoptively transferred OT-I CD8^+^ T cells. Tumor-bearing Thy1.1 mice were infused with activated OT-I CD8^+^ T cells that had been transiently cultured in either methionine-replete (Ctrl ACT) or methionine-restricted medium (transient-MR ACT). Following T cell infusion, mice were maintained on either CD or MRD (**Fig. 5A**). As expected, infusion with OT-I CD8^+^ T cells led to effective tumor control in mice fed CD (**Fig. 5B and 5C**). Transient MR during *in vitro* T cell expansion did not further enhance antitumor activity of OT-I CD8^+^ T cells *in vivo* (**Fig. 5C and 5D**). In contrast, sustained MR through dietary restriction impaired the efficacy of ACT, resulting in accelerated tumor progression (**Fig. 5C and 5F, 5D and 5G, and 5L-5O**) and decreased overall survival (**Fig. 5I and 5J**). Importantly, MRD alone did not alter tumor growth or survival in the absence of ACT (**Fig. 5H and 5K**), indicating that methionine availability specifically influences tumor control mediated by adoptively transferred T cells. Consistent with intratumoral methionine levels, serum methionine concentrations at the endpoint were also significantly lowered in tumor-bearing mice maintained on MRD (Supplementary Fig. S4A), confirming that dietary MR induces systemic methionine restriction. Collectively, these findings demonstrate that while transient MR does not compromise ACT, sustained methionine depletion through dietary restriction disrupts the antitumor function of adoptively transferred T cells, underscoring the requirement for adequate methionine availability to support therapeutic efficacy.

**Figure 5.**
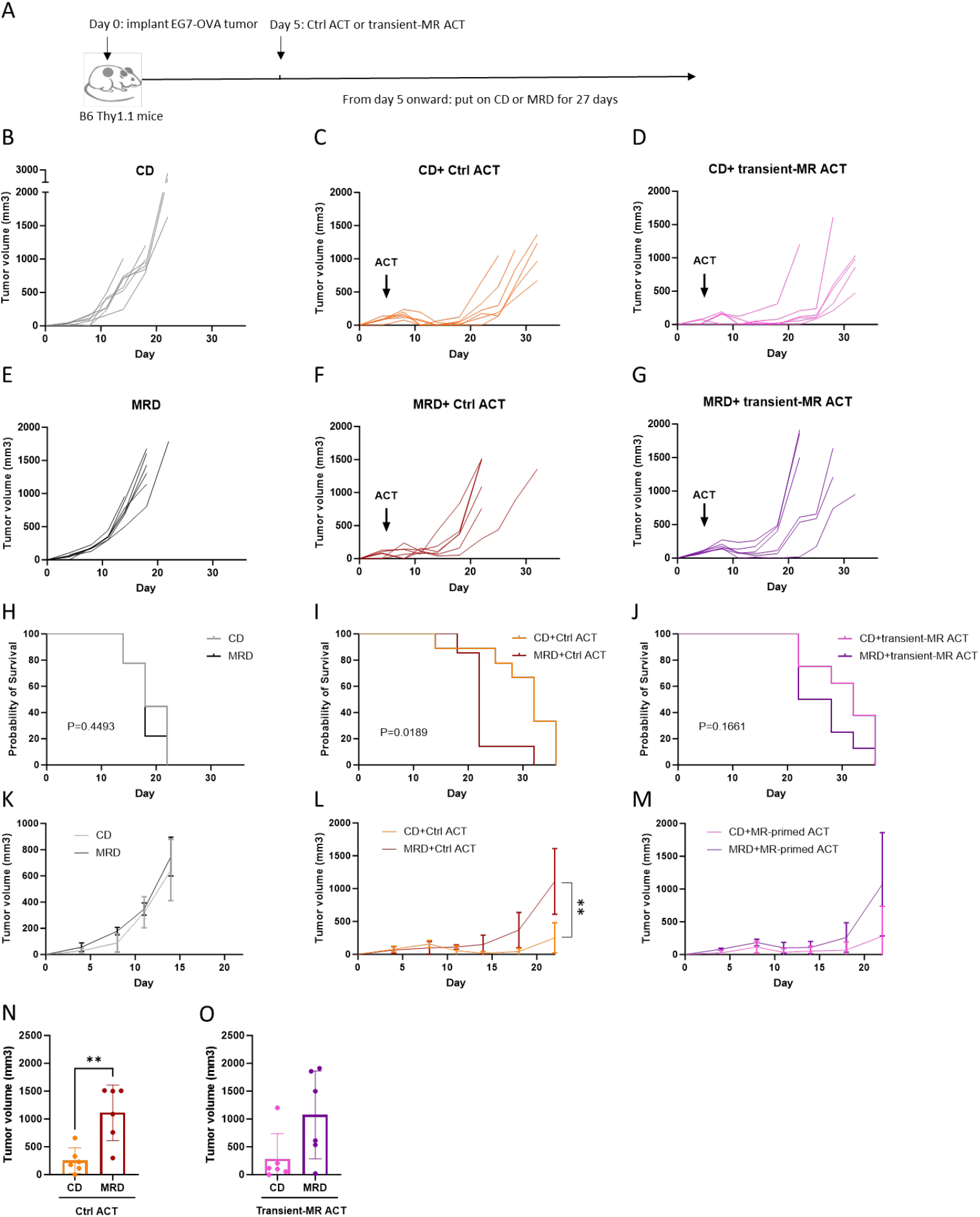
Methionine is required for the antitumor function of adoptively transferred CD8^+^ T cells. **A**, Experimental design. Female B6 Thy1.1 mice (4 months old) were subcutaneously implanted with 1×10^6^ EG7-OVA tumor cells on the right flank (day 0). On day 5, the mice were infused with 6.5×10^5^ activated OT-I CD8^+^ T cells cultured for 16 hours in either control medium (100 μM methionine; Ctrl ACT) or methionine-restricted medium (5 μM methionine; transient-MR ACT). Mice were then placed on either a control diet (CD; 0.86% methionine) or a methionine-restricted diet (MRD; 0.06% methionine) for 27 days. **B**, Individual tumor growth curves of EG7-OVA tumors under the following conditions: control diet only (CD) (n=7, biological replicates). **C**, 100 μM methionine (Ctrl ACT) and on control diet (CD) (n=6, biological replicates). **D**, 5 μM methionine for 16h (transient-MR ACT) and control diet (CD) (n=6, biological replicates). **E**, MRD only (n=7, biological replicates). **F**, 100 μM methionine (Ctrl ACT) and on MRD (n=6, biological replicates). **G**, 5 μM methionine for 16h (transient-MR ACT) and on MRD (n=6, biological replicates). **H-J**, Kaplan-Meier survival curves of indicated treatment groups (H: n=7; I and J: n=6; biological replicates). **K-M**, Tumor growth curves showing average tumor growth of each group (K: n=7; L and M: n=6; biological replicates; L: statistical significance was calculated based on tumor volumes on day 22). **N and O**, Average tumor volumes of indicated treatment groups on day 22 (n=6, biological replicates). **H-O**, Control diet: CD, Ctrl ACT: infusion of OT-I T cells cultured in media containing 100 μM methionine, transient-MR ACT: infusion of OT-I T cells cultured in media containing 5 μM methionine for 16 h. p values were determined using Kaplan–Meier estimate (**H-J**) or unpaired Student’s t-test (**L and N**), ** p ≤ 0.01; error bars represent standard deviation (SD).

## Discussion

Methionine supports T cell activation and function by fueling protein synthesis, redox regulation, nucleotide biosynthesis, and epigenetic remodeling via the generation of SAM through the methionine cycle. While previous studies show that MR enhances endogenous antitumor immunity with immune checkpoint blockade (ICB) (19,20), the role of methionine in ACT remains largely unexplored. This study provides evidence that methionine availability is essential for the antitumor function of adoptively transferred CD8^+^ T cells and highlights a time-dependent relationship between methionine metabolism and T cell function.

Here, we demonstrate that transient MR enhances IFNγ production and cytolytic activity in activated CD8^+^ T cells *in vitro* through suppression of polyamine synthesis. Mechanistically, we show that this effect is mediated through AdoMetDC, which diverts methionine-derived SAM towards polyamine biosynthesis. Both genetic and pharmacologic inhibition of AdoMetDC increased IFNγ production, establishing a functional link between the methionine cycle and polyamine-mediated regulation of T cell effector function. These findings are consistent with prior reports that polyamine synthesis impairs T cell– mediated immunity and can be targeted to improve therapeutic outcomes (33,34).

Under sustained MR, there was induction of exhaustion-associated markers such as PD-1 and TIM-3 in both activated mouse and human T cells. Interestingly, while murine CD8^+^ T cells maintained or even increased IFNγ expression under sustained MR, human T cells exhibited reduced cytokine production, indicating a species-specific sensitivity to methionine availability. This difference may stem from an underlying metabolic disparity, as basal methionine levels are substantially lower in humans (∼23–29 μM) compared to laboratory mice (∼82 μM), making human T cells more susceptible to further methionine restriction. It is also possible that sustained methionine deprivation downregulates methionine transporters such as SLC43A2 (LAT4) or SLC7A5 (LAT1) by decreasing global protein synthesis, reducing cellular methionine uptake. Given that human T cells already function near their methionine threshold due to lower circulating levels, reduced transporter expression could further limit intracellular methionine availability and contribute to their heightened sensitivity to sustained MR.

Given the time-dependent role of methionine in T cell function observed *in vitro*, we investigated whether transient or sustained MR could impact the therapeutic efficacy of adoptively transferred OT-I CD8^+^ T cells in EG7-OVA tumor-bearing mice. Despite the enhancement in T cell function when cultured under transient MR, the beneficial *in vitro* effect did not translate to enhanced antitumor responses *in vivo*. Transient MR during T cell culture did not improve the efficacy of adoptively transferred OT-I CD8^+^ T cells, likely due to re-exposure to physiological methionine levels in the host upon infusion, which may reverse MR-induced metabolic programming.

In contrast, sustained MR through dietary restriction reduced intratumoral methionine levels and led to systemic reduction in methionine levels in tumor-bearing mice. The former was confirmed by two distinct mass spectrometry imaging (MSI) technologies, MALDI and nano-DESI, while the presence of methionine in the MSI-scanned tissues was verified by on-tissue MS/MS (Supplementary Fig. S5 and Fig. S6). These results provide important insights into how intratumoral methionine levels respond to dietary MR, a topic that has not been previously examined despite recent interest in understanding the role of MR in cancer treatment (35). Moreover, sustained MR via dietary restriction significantly impaired the anti-tumor effect of adoptively transferred T cells, leading to reduced tumor control and survival. *In vivo*, MR likely affects other immune cells, as well as stromal cells that support antitumor immunity, and may impair antigen presentation, reduce cytokine production, or weaken co-stimulatory signals, indirectly limiting T cell function. Thus, reduced antitumor activity may stem from broader immunosuppressive effects rather than a T cell–intrinsic defect alone.

Recent work using a metronomic MR strategy, cycling between periods of restriction and repletion, has shown promise in enhancing the antitumor response to ICB while avoiding T cell dysfunction (36). In contrast, our study reveals that sustained MR, particularly in the setting of ACT, leads to reduced therapeutic benefit. These findings reveal a more nuanced role for methionine in T cell biology. While transient MR can temporarily enhance CD8^+^ T cell effector function through suppression of polyamine synthesis, sustained MR induces expression of exhaustion-associated markers in CD8^+^ T cells and impairs antitumor efficacy in the context of ACT.

## Acknowledgments

This study is supported by the Terry Fox New Frontiers Program Project and the Lotte and John Hecht Memorial Foundation Grants (TFRI 1125; JJL), Mitacs Accelerate awards (IT39177; JJL, TZ and IT39215; HP, DRG, MS), Canadian Institutes of Health Research (MOP-142351 and PJT-162279; JJL), IRiCOR, Ovarian Cancer Canada, and the US Department of Defense OCRP (W81XWH-18-1-0264; JJL), University of Victoria Graduate Awards (TZ and MS), Natural Sciences and Engineering Research Council of Canada (MS), Genome British Columbia (GBC) and Genome Canada for The Metabolomics Innovation Centre (TMIC) and the BC Proteomics Centre (365MET, 375MET and 374PRO, DRG). RJD is supported by the Howard Hughes Medical Institute Investigator Program, grant R35CA220449 from the National Cancer Institute, and grant RP240494 from the Cancer Prevention and Research Institute of Texas. Salary support for JM was provided by the Michael Smith Foundation for Health Research (MSHRBC) trainee fellowship (RT-2024-03771), and KDD is supported by funding from the Natural Sciences and Engineering Research Council of Canada (RGPIN-2022-03696), Mitacs Accellerate (IT39215), TFRI PPG 1125, Canadian Foundation for Innovation and BC Knowledge Development Fund (43810), and MSHRBC Scholar program (SCH-2025-04631). JA is funded by Natural Science and Engineering Research Council of Canada (NSERC), Grant RGPIN-2017-04162.

We would like to thank Servier Pharmaceuticals (formerly Agios Pharmaceuticals, Oncology), particularly Dr. Donald Simons, for providing the MAT2A inhibitor AGI-24512. We also thank Dr. Gillian Kingsbury for scientific input. We appreciate the Animal Care Service team at the University of Victoria, especially Dr. Michele Martin, Tracey Sutcliffe, Alison Bohnet, Ahmed Olodo, and Celestine Aniugwu. Additionally, we gratefully acknowledge the invaluable contributions of the animals used in this study.

## Notes

### Competing Interest Statement

RJD is a founder and advisor at Atavistik Bioscience, and an advisor for Faeth Therapeutics and Vida Ventures. The other authors declare no competing financial or non-financial interests related to the content of this communication.

