## Supplementary material for "Methionine regulates antitumor function of CD8^+^ T cells through polyamine synthesis": Suplementry figures, tables, and information

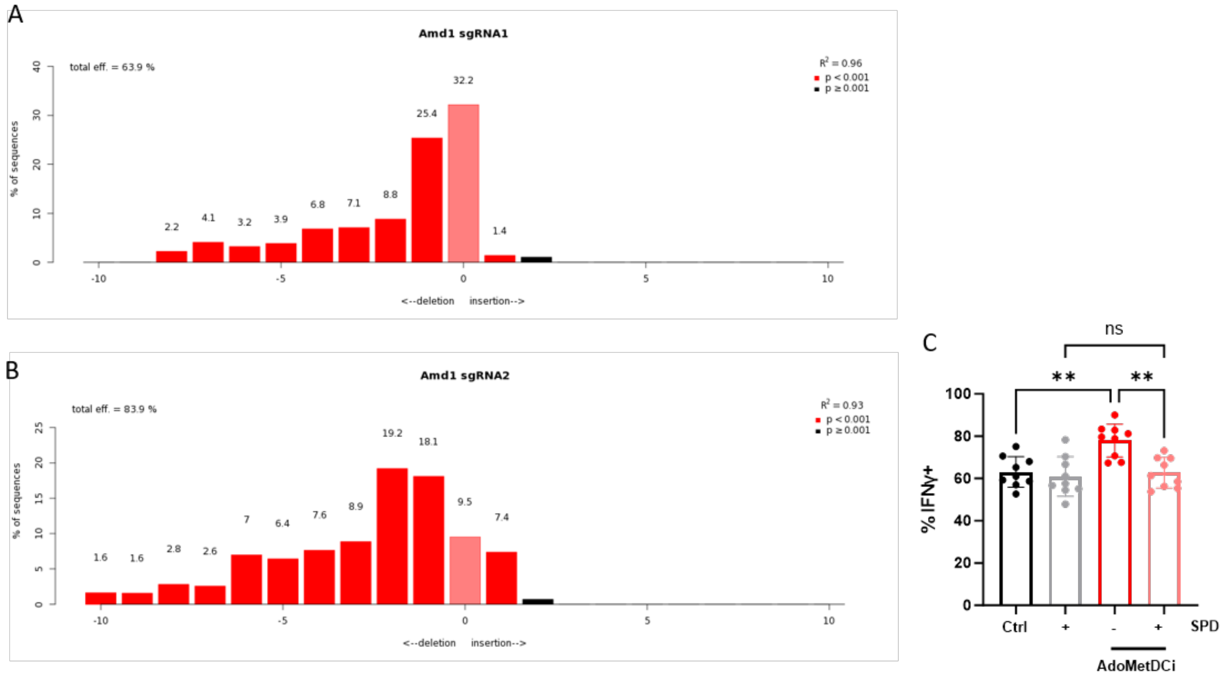


Figure S1. Genetic and pharmacologic inhibition of polyamine synthesis enhances IFNγ expression in CD8⁺ T cells. **A and B**, TIDE analysis showing the indel profiles in Amd1 following CRISPR-Cas9 editing with sgRNA1 and sgRNA2 in mouse CD8⁺ T cells, assessed four days after electroporation. **C**, IFNγ expression in activated mouse CD8⁺ T cells cultured for 40 hours under the following conditions: vehicle control (Ctrl; 0.1% DMSO), AdoMetDC inhibitor (AdoMetDCi; 50 µM CGP-48664), or AdoMetDCi + spermidine (SPD; 30 µM spermidine) (n=9, biological replicates). P-values were determined using one-way ANOVA with Holm-Šídák tests. **p ≤ 0.01. Error bars indicate standard deviation (SD).

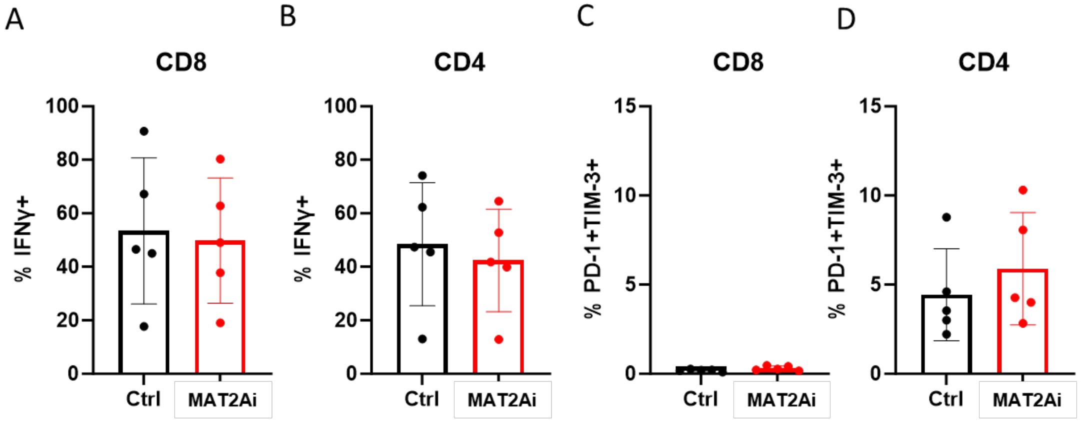


Figure S2. Prolonged inhibition of the methionine cycle induces an exhaustion-like phenotype in human T cells. IFNγ expression in activated human CD8$^{+}$ (**A**) and CD4$^{+}$ (**B**) T cells cultured for 4–5 days in the presence of MAT2A inhibitor (MAT2Ai; 5 µM AGI-24512) or vehicle control (Ctrl; 0.1% DMSO) (n=5, biological replicates). (**C and D**) Expression of exhaustion-associated markers (PD-1$^{+}$/TIM-3$^{+}$) in the same human CD8$^{+}$ (**C**) and CD4$^{+}$ (**D**) T cell populations under the indicated conditions (n=5, biological replicates).

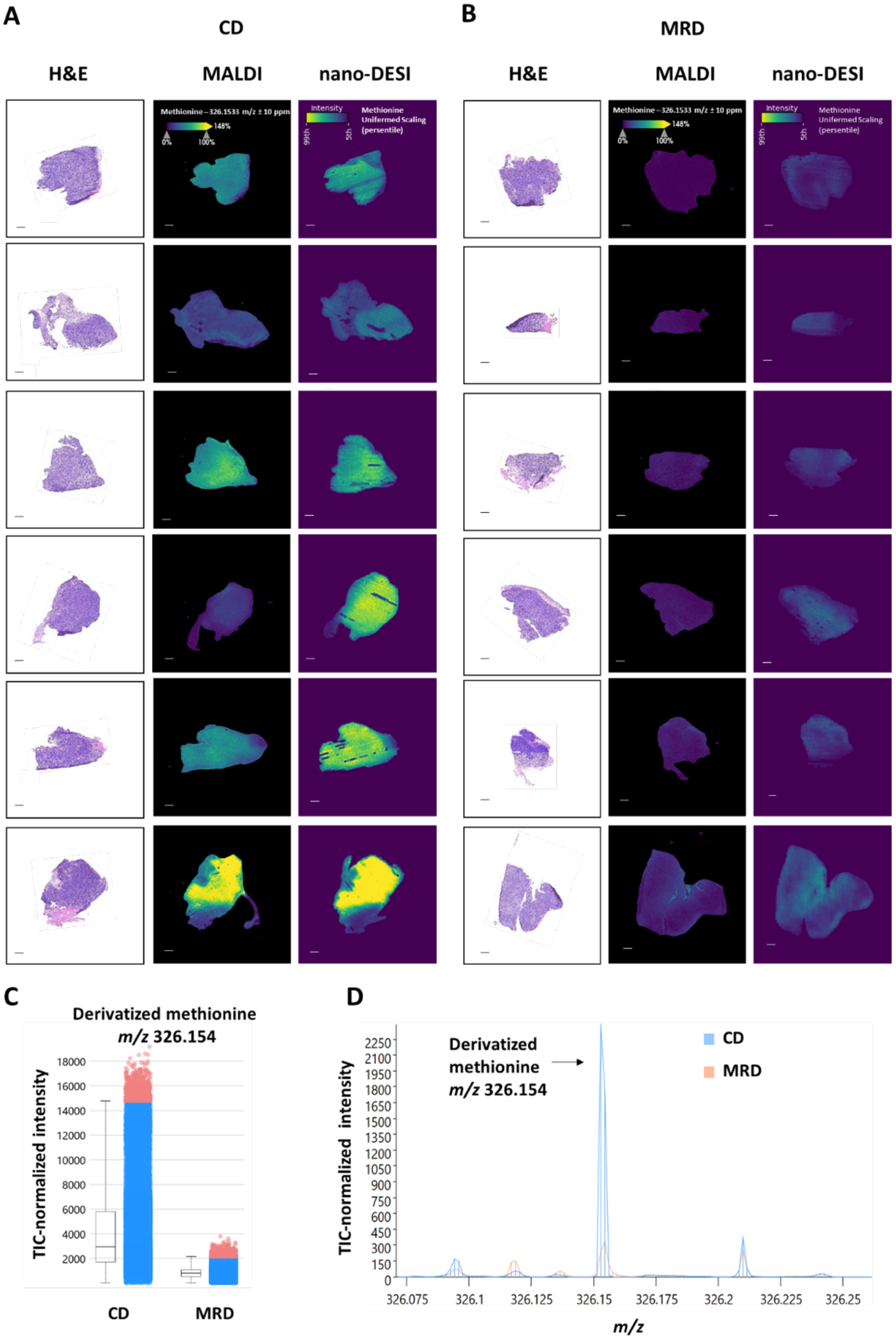


Figure S3. Representative histochemistry and mass spectrometry images of all 12 tumors obtained in the experiment are shown in **Fig. 4. A**, Representative H&E, MALDI, and nano-DESI images of tumor sections from mice placed on CD (n=6, biological replicates). **B,** Representative H&E, MALDI, and nano-DESI images of tumors from mice placed on MRD (n=6, biological replicates). **C**, Box and bar plot showing pooled TIC-normalized intensities of methionine in tumors placed on CD or MRD, obtained by MALDI. Open boxes show intensities from the 25-75$^{th}$ percentile with middle lines showing the median intensity and whiskers indicating the 75-99$^{th}$ percentile (top) and the 0-25$^{th}$ percentile (bottom); Blue bars show intensities form the 0-99$^{th}$ percentile with the red dots corresponding to the 99-100$^{th}$ percentile. **D**, Integrated peak plot showing pooled TIC-normalized intensities of methionine in tumors placed on CD or MRD, obtained by MALDI.

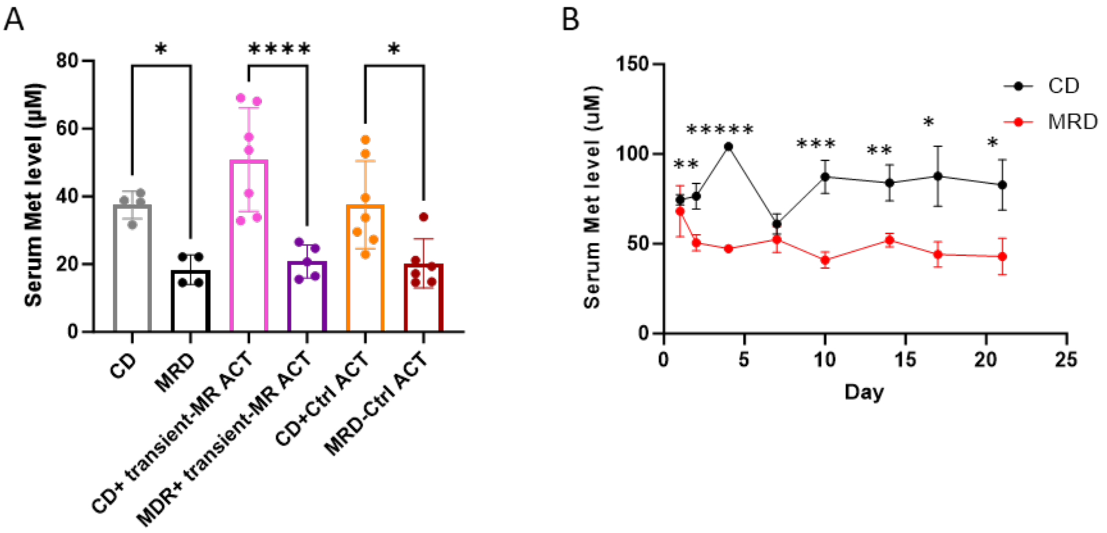


Figure S4. Methionine-restricted diet reduces serum methionine levels in tumor-bearing and healthy mice. **A,** Serum methionine (Met) levels in EG7-OVA tumor-bearing mice received indicated treatments (samples collected in the same experiment shown in Fig. 4). Control diet: CD, Ctrl ACT: infusion of OT-I T cells cultured in media containing 100 µM Met, transient-MR ACT: infusion of OT-I T cells cultured in media containing 5 µM Met for 16 h (n=4-7, biological replicates). **B,** Longitudinal changes in serum methionine (Met) levels in healthy mice fed on the control diet (CD) or MRD over 3 weeks (n=3, biological replicates). P-values were calculated using one-way ANOVA with Holm-Šídák tests (A) or unpaired Student's t-tests (B). $*p \leq 0.05$, $**p \leq 0.01$, $***p \leq 0.001$, $*****p \leq 0.00001$. Error bars represent standard deviation (SD).

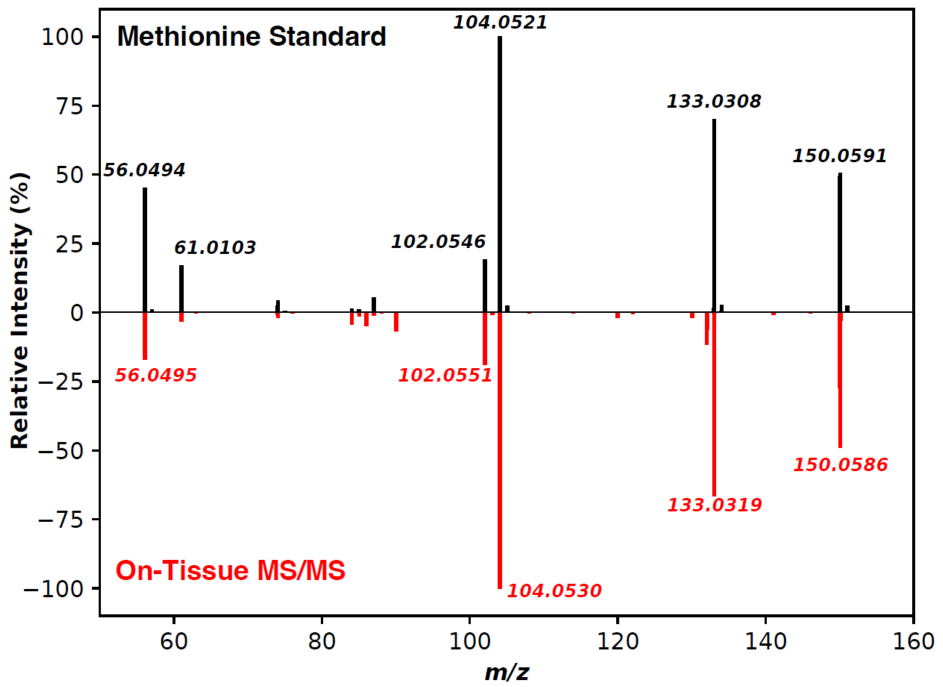


Figure S5. Normalized product ion scan (HCD: 30%) for a methionine standard solution (top; black) and on-tissue MS/MS (bottom; red) confirming the presence of methionine in the tissue. On-tissue MS/MS was collected from a nano-DESI-imaged tumor section shown in Fig. 5, averaged over a full line-scan of the tissue.

**A**

Methionine standard

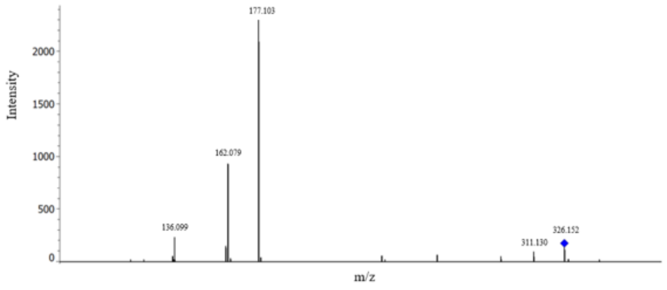


**B**

CD tumors

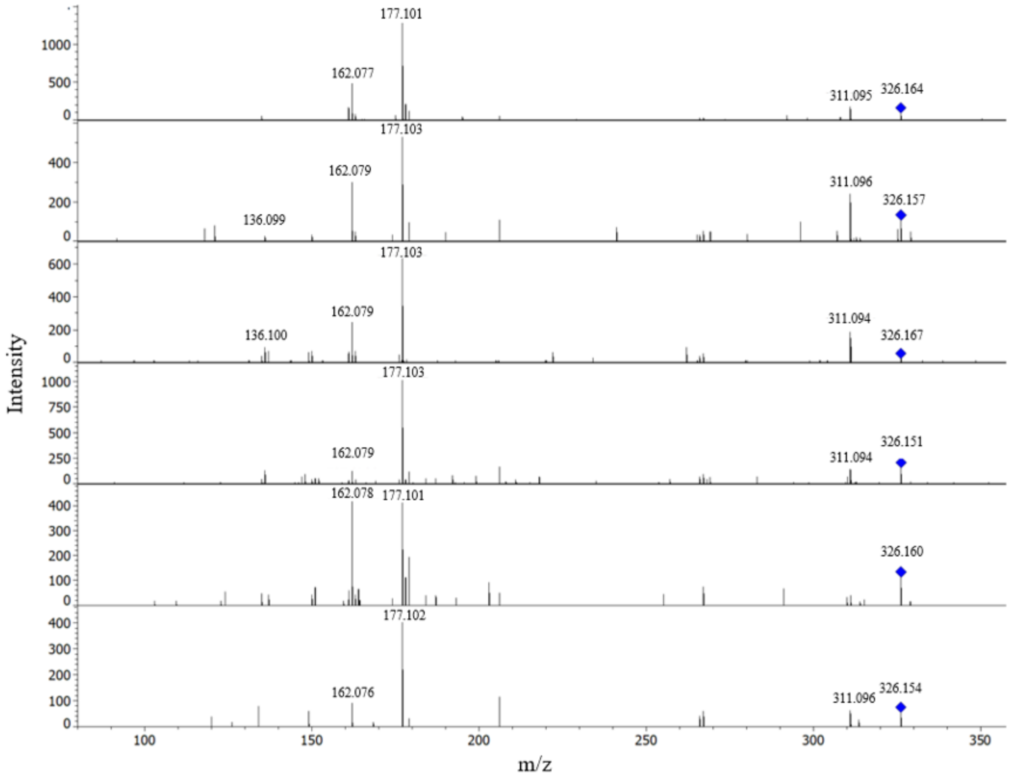

C

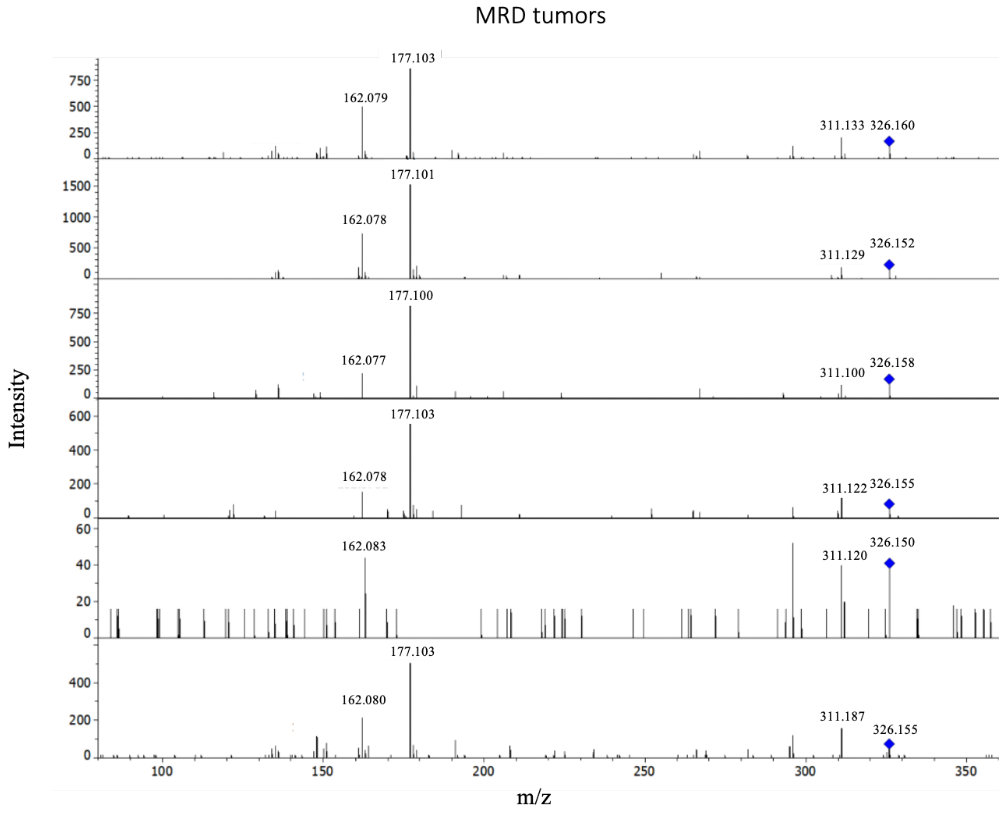


Figure S6. On-tissue MS/MS fragmentation spectra of a spotted methionine standard (**A**), CD and MRD tissues (**B**, **C**), confirming the presence of methionine in tumor sections imaged by MALDI shown in Fig. 5. After MALDI-MSI, tissues were subjected to on-tissue MS/MS with a precursor ion *m/z* 326.154 representing the TAHS derivative of methionine. Collision energy was increased to 25 eV until fragment ion masses were observed. Tandem mass spectra from each tissue were analyzed using Data Analysis software (Bruker) and proposed structures were generated using the Fragmentation Explore tool.

Table S1. Nutrient inclusion rate of the control diet (CD) and the methionine-restricted diet (MRD)

| Nutrient | CD (g/kg) | MRD (g/kg) |
|---|---|---|
| L-Alanine | 3.5 | 3.5 |
| L-Arginine HCl | 12.1 | 12.1 |
| L-Asparagine | 6 | 6 |
| L-Aspartic Acid | 3.5 | 3.5 |
| L-Cystine | 10 | 10 |
| L-Glutamic Acid | 40 | 40 |
| Glycine | 23.3 | 23.3 |
| L-Histidine HCl, monohydrate | 4.5 | 4.5 |
| L-Isoleucine | 8 | 8 |
| L-Leucine | 12 | 12 |
| L-Lysine HCl | 18 | 18 |
| **L-Methionine** | **8.6** | **0.6** |
| L-Phenylalanine | 7.5 | 7.5 |
| L-Proline | 3.5 | 3.5 |
| L-Serine | 3.5 | 3.5 |
| L-Threonine | 8.2 | 8.2 |
| L-Tryptophan | 1.8 | 1.8 |
| L-Tyrosine | 5 | 5 |
| L-Valine | 8 | 8 |
| Sucrose | 123.5 | 123.5 |
| Corn Starch | 374.23 | 382.255 |
| Maltodextrin | 150 | 150 |
| Soybean Oil | 50 | 50 |
| Cellulose | 50 | 50 |
| Mineral Mix, AIN-93M-MX (94049) | 35 | 35 |
| Calcium Phosphate, monobasic, monohydrate | 8.2 | 8.2 |
| Vitamin Mix, AIN-93-VX (94047) | 19.5 | 19.5 |
| Choline Bitartrate | 2.5 | 2.5 |
| TBHQ, antioxidant | 0.02 | 0.02 |
| Orange Food Color | 0.05 | 0.05 |

Table S2. Sprayer parameters for MALDI MSI

| HTX M5 Spray Settings for TAHS | |
|---|---|
| Derivatization agent | TAHS |
| Concentration (mg/mL) | 1.25 |
| Solvent | ACN |
| Density on slide (mg/mm^2) | 2.08 e-4 |
| Nozzle temp (°C) | 60 |
| Plate temp (°C) | 30 |
| # Passes | 6 |
| Flow rate (µL/min) | 100 |
| Velocity (mm/min) | 1200 |
| Track spacing (mm) | 3 |
| Pattern | CC |
| Pressure (psi) | 10 |
| Gas flow rate (L/min) | 2 |
| Dry time (s) | 0 |
| Nozzle height (mm) | 40 |

Table S3. Sprayer parameters for MALDI MSI

| HTX M5 Spray Settings for 2,5-DHB | |
|---|---|
| Derivatization agent | 2,5-DHB |
| Concentration (mg/mL) | 15 |
| Solvent | 90% ACN + 0.1% TFA |
| Density on slide (mg/mm^2) | 7.20 e-3 |
| Nozzle temp (°C) | 60 |
| Plate temp (°C) | 30 |
| # Passes | 14 |
| Flow rate (µL/min) | 125 |
| Velocity (mm/min) | 1200 |
| Track spacing (mm) | 3 |
| Pattern | CC |
| Pressure (psi) | 10 |
| Gas flow rate (L/min) | 3 |
| Dry time (s) | 0 |
| Nozzle height (mm) | 40 |

Supplementary Information. Code for generating the violin plot of pooled methionine intensities in tumors (Fig. 4C), using Matplotlib 3.10.3:

```
from viu_chem import MSI_Process as msi
import numpy as np
import matplotlib.pyplot as plt
import os
import matplotlib
import scipy.stats

# Setup plotting library
matplotlib.rc('font', family='sans-serif')
matplotlib.rc('font', serif='Helvetica Neue')
matplotlib.rcParams['pdf.fonttype'] = 42 #Allows editable text in Adobe Illustrator
matplotlib.rcParams['ps.fonttype'] = 42 #Allows editable text in Adobe Illustrator

# Define which codes go with which treatment
MR_codes = ["TF26M", "TF27N", "TF28M", "TF32Q", "TF33Q", "TF34I"]
control_codes = ["TF23R", "TF24M", "TF25M", "TF29I", "TF30M", "TF31M"]

# Define where the data and region of interest maps are stored (directories containing all files)
data_folder = "../../LUM2 Sample Storage/VIU Data/Polylysine - SCiLS Compatible"
ROI_folder = "../../LUM2 Sample Storage/VIU Data/ROI_select"

filter_string = "_FTMS + p ESI Full ms [70.imzML" #Filter string that all the imzML files will terminate with

# Function to find the correct files to extract data from
def find_datafiles(code:str,dir:str=data_folder,roi_path:str = ROI_folder, pattern:str=filter_string):
    avail_files = os.listdir(dir)
```

```
    tgt_folder = [file for file in avail_files if code in file][0] #Grabs the corresponding sample code from the target directory
    tgt_file = msi.find_data_filt_string(os.path.join(dir,tgt_folder),pattern) #Finds the right imzML file for that code

    avail_roi = os.listdir(roi_path)
    tgt_roi = [roi for roi in avail_roi if code in roi][0] #Finds the matching roi
    full_roi = os.path.join(roi_path,tgt_roi)

    return tgt_file, full_roi

#Function to extract the flattened image matrix
def get_masked_image(path, roi_path, mz:float, norm:float="tic",ret:str ="cleanup",tolerance:float=10):
    data = msi.get_image_matrix(path,mz,tol=tolerance) #Retrieves matrix of xy pixel intensities as an np array
    if norm == "tic":
        tic = msi.get_TIC_image(path)
        data = np.divide(data, tic)
    elif norm == "wm":
        wm = msi.get_weighted_median_image(path)
        data = np.divide(data,wm)
    elif norm == "combo":
        int_std = msi.get_image_matrix(path, 151.05591)
        tic = msi.get_TIC_image(path)
        data = np.divide(data,int_std, out=np.zeros_like(data),where=int_std!=0)
        data = np.divide(data, tic)
    else:
        int_std = msi.get_image_matrix(path, norm)
        data = np.divide(data,int_std, out=np.zeros_like(data),where=int_std!=0)

    roi = np.load(roi_path)['roi_mask']
```

```
    masked = np.where(roi==1, data,np.nan)
    if ret == "image":
        return masked

    cleanup = masked.flatten()
    cleanup = cleanup[~np.isnan(cleanup)]
    return cleanup


#Function to draw the plots
def draw_violin_plot(analyte:tuple=("Methionine", 150.0583),
norm_method:str="TIC",norm_pass='tic',norm_limits:tuple=None,savepath:str=None, tol:float=3):
    tgt_name, tgt_mz = analyte

    data = {
        "control": [],
        "met_res": [],
    }

    ind_controls = []
    ind_met = []

    # Iterate through each file and add it to the appropriate data category
    for idx, (control, met_res) in enumerate(zip(control_codes,MR_codes)):
        print(f"Starting idx {idx+1} of {len(control_codes)}")
        control_path, control_roi = find_datafiles(control)
        tmp = get_masked_image(control_path, control_roi,tgt_mz,norm=norm_pass, tolerance=tol)
        data["control"].extend(tmp)
        ind_controls.append(np.mean(tmp))
```

```
        met_res_path, met_res_roi = find_datafiles(met_res)
        tmp = get_masked_image(met_res_path, met_res_roi,tgt_mz,norm=norm_pass, tolerance=tol)
        data["met_res"].extend(tmp)
        ind_met.append(np.mean(tmp))


    #Plots the comparison, controls on the left (low) side, MR on the right (high) side
    plt.violinplot(data['control'],side='low',showextrema=False,points=1000)
    plt.violinplot(data['met_res'],side='high',showextrema=False,points=1000)
    plt.ylabel(f"{tgt_name} / {norm_method}")
    plt.title(f"Global distribution of {tgt_name} - {norm_method} normalized")
    plt.xticks([1],labels=["CD | MRD"])
    if norm_limits:
        plt.ylim(norm_limits)
    plt.tight_layout()
    if savepath:
        plt.savefig(savepath)
    plt.show()

    return data['control'], data['met_res'], ind_controls, ind_met

def run_stats(data):
    controls, mr_data, ind_controls, ind_met = data
    #Computes the t-statistic/pval for the entire pixel population (tissue only)
    result = scipy.stats.ttest_ind(controls, mr_data)
    print(f"t-stat: {result.statistic}\np-val: {result.pvalue:.12f}\n degrees of freedom: {result.df}")

    #Computes the t-statistic/pval for mean signal of each image (tissue only)
    result2 = scipy.stats.ttest_ind(ind_controls,ind_met)
    print(f"t-stat: {result2.statistic}\np-val: {result2.pvalue:.12f}\n degrees of freedom: {result2.df}")
```

```
if __name__ == "__main__":
    norm_method = "TIC"
    norm_pass = 'tic'
    ylims = [0, 0.012]

    data = draw_violin_plot(analyte=("Methionine", 150.0583),tol=3,norm_method=norm_method,
norm_pass=norm_pass, norm_limits=ylims)
    run_stats(data)
```
